# Study on the efficacy of *Cynodon dactylon* against Infectious Myonecrosis Virus (IMNV) in *Penaeus vannamei*

**DOI:** 10.1101/2020.11.09.374074

**Authors:** Rajeev Kumar Jha, Soy Daniel Wisoyo, Kristina, Sarayut Srisombat

**Affiliations:** PT. Central Proteina Prima, Indonesia

**Keywords:** IMNV, *Cynodon dactylon* plant extract. survival rate

## Abstract

Infectious myonecrosis virus (IMNV) is one of the most pathogenic viruses causing severe mortality in *Penaeus vannamei* in many countries. Several strategies have been implemented to inhibit the presence of IMNV disease. The present study was carried out to examine the antiviral activity of *Cynodon dactylon* ethanolic extract on IMNV disease in shrimp by *in vivo* testing. The *C. dactylon* plant extract was incorporated with artificial pellet feed at a concentration of 10, 15%, 20%, and 25%. For the experimental challenge shrimp were fed with IMNV-infected shrimp meat. The antiviral activity was determined by observing survival rates, and IMNV infection was confirmed at the end of the experiment through polymerase chain reaction (PCR) identification. This study showed that the plant extract of *C. dactylon* was found to be highly effective in preventing IMNV infection with up to 98% survival rate in *Penaeus vannamei*.

## INTRODUCTION

The shrimp, *Penaeus vannamei*, is one of the main species for aquaculture in many countries. It Indonesia alone, it has been cultured in at least 17 provinces (East Java, Central Java, West Java, Yogyakarta, Banten, Bali, West Nusa Tenggara, Lampung, South Sumatra, Riau, Bengkulu, West Sumatra, North Sumatra, South Sulawesi, South Kalimantan, East Kalimantan, and West Kalimantan) (The et al., 2010). There were reports from Indonesian shrimp farmers of high mortality in cultivated *P. vannamei* with gross signs of white muscle in 2006 (Senapin *et al*., 2007). Similar symptoms also found in several countries like north-eastern Brazil (Andrade *et al*., 2007; Lightner *et al*., 2004, Nunes *et al*., 2004; Poulos *et al*., 2006, Prasad *et al*., 2017) as well as in other South-East Asian countries. A study on the infected shrimp revealed the presence of IMNV using a standard commercial detection kit (Senapin *et al*., 2007). Infectious myonecrosis virus (IMNV) will present focal to extensive white necrotic areas in striated (skeletal) muscles, especially in the distal abdominal segments and tail fan, which can become necrotic and reddened in some individual shrimp. Severely affected shrimp become moribund, and mortalities can be instantaneously high and continue for several days (Senapin *et al*., 2007).

Several strategies have been continuously carried out to prevent the spread of IMNV infection, among the good management practice of shrimp farming, specific pathogen-free brood stock, and disinfection of eggs and larvae. However, to control the viral spread, the limited knowledge of the mode of action and various agents’ application restricts the success. One of the most promising methods of preventing diseases is strengthening the shrimp defense mechanisms through immunostimulant administration (Balasubramanian *et al*., 2008; Kaleeswaran *et al*., 2011). Immunostimulants are enhancing both specific and nonspecific immunity against infectious diseases. Various studies have been carried out to obtain immunostimulants’ performance to improve immune response and reduce disease impacts. Jha et al. 2016, has successfully developed a formulation of essential oil blend extracted from ten plants as a feed additive and determine the efficacy against WSSV.

*Cynodon dactylon* is widely available and distributed in almost all parts of the world. This herbal plant is known for its ability to treat hysteria, epilepsy, and insanity. *C. dactylon* is used for purifying the blood, anuria, biliousness, conjunctivitis, diarrhea, gonorrhea, itches, and stomachache (Kaleeswaran *et al*., 2011). The aqueous and ethanolic extracts of *C. dactylon* have shown antiviral properties against WSSV for *Penaeus monodon* (Balasubramanian *et al*., 2008). A trial was conducted to determine the efficacy of shrimp feed supplemented with *C. dactylon* ethanolic extract against IMNV in *Penaeus vannamei*.

## MATERIALS AND METHODS

### Preparation of Plant Extract from *C. dactylon*

*C. dactylon* (whole plant except root) was collected from the fields and cleaned. The fresh specimen was then shade dried and powdered using an electrical blender. The powdered plant ware mixed with fresh water in a 1:10 ratio and heated at 60^0^C for 20-30 minutes. A crude yellow aqueous extract was obtained by filtering out the residues, and the filtrate was diluted with absolute ethanol in the ratio of 1:4. A copious quantity of gel-like precipitate formed immediately after the addition of ethanol. The supernatant of the ethanol extract was recovered by filtration, and the gel pellet was discarded. The supernatant solution was subjected to distillation at 60^0^C for 2-3 days to obtain a plant extract. The obtained plant extract was completely air-dried. The dried product was sealed in a glass container and kept in the refrigerator for further use.

### Preparation of *C. dactylon-*Supplemented Feed

The plant extract from *C. dactylon* was incorporated with commercially available artificial pellet feed (CP Prima, Indonesia) at the concentration of 10%, 15%, 20%, and 25% inclusion. The required concentration of plant extract was dissolved in the water, sprayed on the commercial feed on ice, and incubated for 30 min to absorb the plant extract. Then, it was re-coated with squid oil to prevent the plant’s dispersion extract in water. The feed was then dried and stored at room temperature until given to shrimp.

### Investigation of anti-IMNV activity of *C. dactylon*

#### a. IMN Virus Preparation for Injection inoculation

The IMNV-infected shrimp were collected and checked for the presence of the virus. The head, shell, and gut were removed, and the muscle was first to cut into small pieces with the help of scissors and then minced into fine pieces by using a sterilized manual blender. The obtained semi-solid gel-like muscle was filtered through an 80-micron filter and recovered. The obtained minced muscle carrying IMNV was homogenized and checked for the presence and quantification by RT-PCR. The minced muscle with IMNV was packed in small Blocks stored at −20^0^C before use. At the time of using were thawed slowly shifting to cooler temperature (−20^0^C to 4^0^C and finally to 18-20-25^0^C. The complete procedure took 4-5 hours of time.

#### b. Laboratory Experiment of IMNV-challenged Shrimp

Shrimps were collected from Marine Research Center, Lampung. The treatment tanks were divided into four groups of 20 shrimp per tank. The trial duration was 27 days, including three days of acclimatization, 14 days of feeding; at day 15, shrimp were challenged with IMNV. From day 16 onwards, it continues feeding and observation for ten days. The method in detail was as followed:

##### c. Tank and water preparation

The transparent plastic tanks with 125 L capacity are used for the lab-scale trial. The marine chlorinated water was obtained from CPB Hatchery. The water temperature was maintained at 25±1^0^C. The water salinity was maintained at 20 ppt, pH 7.6-8, Dissolved Oxygen 5.1-6 ppm, the water exchange was done at the rate of 20% daily. Siphoning was done once a day.

##### Treatment groups

In negative control group, shrimp were fed with normal pellet feed on regular CP-3 feed throughout the experimental period. In positive control group, the shrimp were fed with normal pellet feed and challenged with 1.8% IMNV muscle on day 15 and continued feeding with regular feed from day 16 onwards. In each treatment group, shrimp were fed with *C. dactylon* extract supplemented feed (RAV-S) for the first 14 days and challenged with IMNV at day 15 and feeding continued throughout the experimental period.

##### Challenge method

Shrimps was challenged per *os* with 1% IMNV-infected muscle.

##### Post challenge observation

The challenged shrimp were under intense observation for their behavior and feeding rate and cumulative mortality during the trial. The feeding rate and tank bottom siphoning schedule and maintained the same for both control and treatment group throughout the experiment. Uneaten food and waste matter were removed before feeding. The experimental animals were examined twice a day for gross signs of disease and the number of deaths was recorded. The dead and the moribund shrimp were taken out and confirmed by

#### d. Cage Scale Trial of IMNV-challenged Shrimp

The shrimp were divided into 5 groups of 40 shrimp per cage and each group was conducted in three replicates. The duration of trial was of 35 days including 10 days of acclimatization, 14 days feeding, at day 15 shrimp were challenged with IMNV and from day 16 onwards continues feeding and observation for 10 days. The method in detail were as followed,

##### Cage and water preparation

The water preparation was done by following the standard procedure of culture pond preparation. The 3 cages each group (1 m^3^ size and 4 mm mesh size) were set in 150 m^2^ raceways. The water depth maintained at least at 70 cm and maximum to 1 m. The marine chlorinated water was obtained from CPB Hatchery. The water temperature was maintained at 25±1^0^C. The water salinity was maintained at 20 ppt, pH 7.6-8, Dissolved Oxygen 5.1-6 ppm, no water exchange during the experimental period.

##### Treatment groups

Feeding treatment in the cage scale experiment was carried out the same as on the lab scale. Specific Pathogen Free (SPF) shrimp were stocked with stocking density of 80/m^2^ with shrimp average body weight is 4 g. Acclimatization was carried out for 2-3 days and started feeding.

##### Challenge method

Shrimps was challenged per *os* with 2% IMNV-infected muscle. Day 16 onwards continued feeding and mortality and behavior were observed.

##### Post challenge observation

Observation on the challenged shrimp were done as the lab scale experiment. The water quality parameters checked on daily basis (two times normally). The total Bacterial Count, Total *Vibrio* Count, plankton population and virus (WSSV and IMNV) would be checked in every five days. The data of full moon, new moon, daily rain fall were also be obtained and recorded. The data of full moon, new moon, daily rain fall were also be obtained and recorded.

#### e. Cage-Pond Scale Experiment

Cage-Pond trial were conducted in commercial ponds in IMNV prone area. The 3×1 m cage were prepared following the standard and water preparation adopted from CPB’s SOP. Water for experiment were treated with 1 ppm of CuSO_4,_ 1 ppm of Pondfos, and 20 ppm of Chlorine. The SPF shrimp were stocked with the same density of 80 pieces/m^2^ in each cage. Shrimp were cultured in autotrophic system with less water exchange and high aeration. Water was maintained at maximum level (120 cm) during the culture and was added only to replace the loose water by siphoning, leaking and evaporation. The treatment group were fed with *C. dactylon-*supplemented feed (RAV-S) that were prepared in CP Feed mill in Surabaya, whereas the control group were fed with CP regular feed.

## RESULTS and DISCUSSIONS

The occurrence of IMNV infection in *P. vannamei* in Indonesia was confirmed after sequencing the PCR fragment of IMNV from shrimp sample originated from East Java in 2006 (Senapin *et al*., 2007). Subsequent analysis revealed that the Indonesian IMNV sample had 99.6% nucleic acid sequence identity than the Brazilian IMNV reported at GenBank (Nur, 2007; Senapin *et al*., 2007). This disease affects the juvenile and grows out shrimp (DOC 30 - 90) with 10-30 % mortalities. The outbreak did not appear to be season dependent (Nur, 2007). The development of antiviral plant extract has been done against WSSV to control the spreading of this disease and protect cultured shrimp from this virus. In our previous work, the application of *Cynodon dactylon* extract has anti-WSSV properties that protect shrimp from WSSV infections. In the present study, an attempt was made to explore the efficacy of the administration of *C. dactylon* (RAV-S) to protect shrimp against IMNV.

Anti-IMNV activity of *C. dactylon* was determined by observing the survival of shrimp through an *in vivo* challenge test both in a lab-scale experiment and in a cage scale experiment. From the lab-scale experiment, complete mortality was observed in positive control within four days after infection. Ten days after infection, the cumulative mortality was found significantly higher in positive control (100%). Mortality in treatment group was observed from day 4 post-infection. The highest protection against IMNV is the treatment group with 25% and 20% of *Cynodon* extract (30% protection) (Figure 1), followed by the treatment group with 10% and 15% of *Cynodon* extract, respectively.

**Figure 1.**
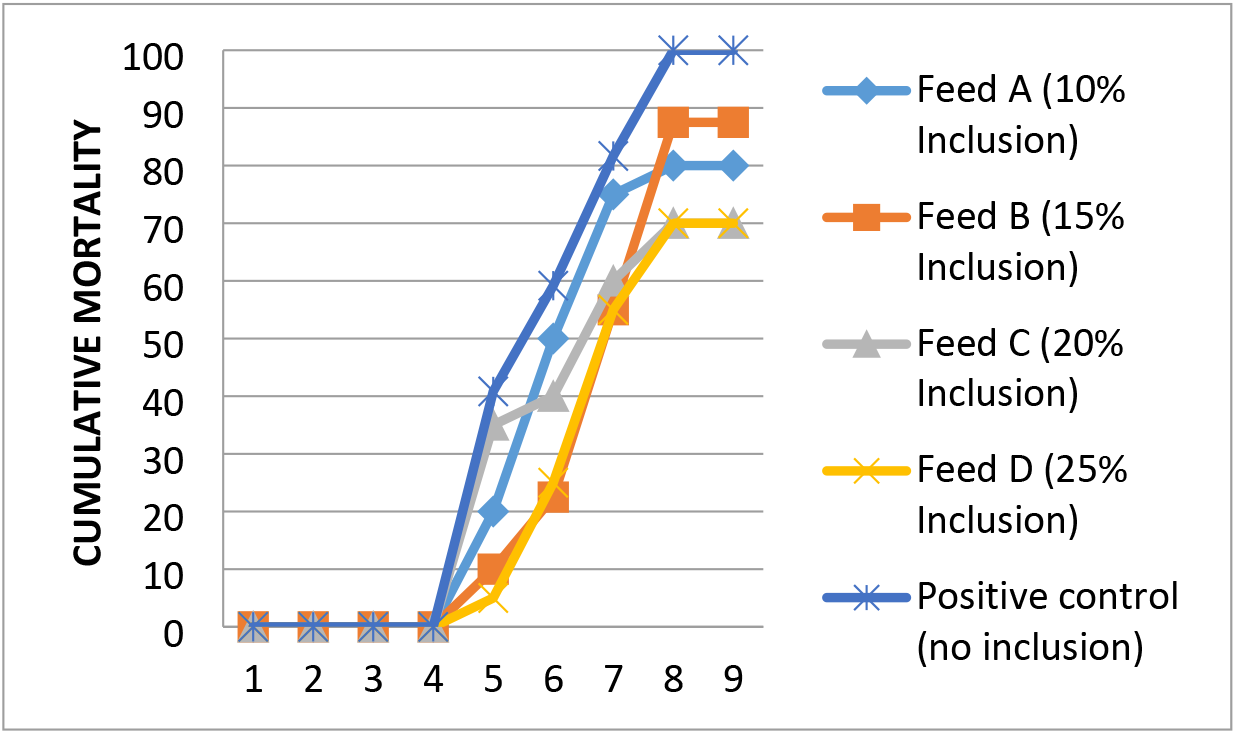
Percentage of cumulative mortality of *L.vannamei* in the *in vivo* challenge test with IMNV and four different concentrations (10, 15, 20 and 25 mg/kg) of *C. dactylon* crude extract

The cage scale experiment was conducted to re-confirm the efficacy of *Cynodon dactylon* (RAV-S feed) to control IMNV infection in shrimp culture. Effect of *Cynodon dactylon*-coated feed on shrimp survival rate found satisfactory. The difference in survival was clear from each treatment (Table 1), with the highest protection provided by feed C (SR 91.1%) after 60 days post infection and followed by feed B with SR 75.2% but the positive control also had a high survival (79.6%). The major difference between survived positive control and *C. dactylon* feed were observed in presence of clinical sign. In the positive control group, the percentage of IMNV signs was 50%, it was higher than the Feed B group (28.9%) and Feed C (2.55%). This could be concluded that *C. dactylon* has the ability to protect shrimp from IMNV and reduce the appearance of disease symptoms.

**Table 1.** Percentage of Shrimp Cumulative Mortality from Cage Experiment

| Treatment | Cage | shrimps (n) | Mortality | Healthy Shrimps | %SR |  | Total Gross sign |  |  |  |  |  |  |  |  |  |  |
| --- | --- | --- | --- | --- | --- | --- | --- | --- | --- | --- | --- | --- | --- | --- | --- | --- | --- |
|  |  |  |  |  | SR/cage | Ave | G-01 |  | G-02 |  | G-03 |  | G-04 |  | All |  | Ave |
| Feed B | 1 | 86 | 24 | 62 | 72.09 | 75.2 | 4 | 5.90% | 10 | 14.70% | 2 | 2.90% | 1 | 1.50% | 17 | 25% | 28.90% |
|  | 2 | 86 | 23 | 63 | 73.26 |  | 6 | 8.50% | 15 | 21.10% | 1 | 1.40% | 0 | 0% | 22 | 30.90% |  |
|  | 3 | 86 | 17 | 69 | 80.23 |  | 4 | 6.20% | 15 | 23.10% | 1 | 1.50% | 0 | 0% | 20 | 30.80% |  |
| Feed C | 1 | 45 | 3 | 42 | 93.33 | 91.1 | 1 | 2.60% | 0 | 0% | 0 | 0% | 0 | 0% | 1 | 2.60% | 2.55% |
|  | 2 | 45 | 5 | 40 | 88.89 |  | 0 | 0% | 1 | 2.50% | 0 | 0% | 0 | 0% | 1 | 2.50% |  |
| Positive Control | 1 | 79 | 21 | 58 | 73.42 | 79.6 | 4 | 7.40% | 23 | 42.60% | 0 | 0% | 0 | 0% | 27 | 50% | 50% |
|  | 2 | 77 | 16 | 61 | 79.22 |  | 5 | 12.50% | 18 | 45% | 0 | 0% | 0 | 0% | 23 | 57% |  |
|  | 3 | 80 | 11 | 69 | 86.25 |  | 2 | 3.50% | 23 | 40.40% | 0 | 0% | 0 | 0% | 25 | 43% |  |
| Negative control | 1 | 80 | 0 | 80 | 100 | 99.17 | No Gross Sign |  |  |  |  |  |  |  |  |  |  |
|  | 2 | 80 | 0 | 80 | 100 |  |  |  |  |  |  |  |  |  |  |  |  |
|  | 3 | 80 | 2 | 78 | 97.5 |  |  |  |  |  |  |  |  |  |  |  |  |

The water quality parameters were checked on daily basis (two times a day) and the total plankton were checked every five days. The results showed that there is no significant difference between treatments and control groups in all parameters. It shows that *C. dactylon* crude extract not affecting the water quality and the plankton load.

The cage-pond experiment was carried out to reassure the efficacy of *Cynodon dactylon* (RAV-S feed) to control IMNV infection in shrimp culture. The results of the experiment showed that shrimp given *C. dactylon* crude extract feed had a higher survival rate compared to the control, this was also supported by lower gross sign data compared to the control group.

| Cage (sanjivani feed ) | Pond (reguler feed) 01:41:14 |
| --- | --- |
| SR : 69 % | SR : 53 % |
| Gross sign : 10% | Gross sign: 46% |

The results from this study were found to be encouraging in IMNV-challenged from all the experiments. We observed that shrimp fed with RAV-S-C (20% inclusion) provides 91.1% protection of *L. vannamei* with 2% of the clinical signs of IMNV up to 60 dpi. The results indicate that the extracted chemicals of *C. dactylon* might contain anti-IMNV properties either independently or combinedly and those chemicals can provide complete protection of the animals against the IMNV. However, the concentration of 15% might protect the shrimp by up to 78% during the challenge test period. From this result, it can be concluded that those chemicals substances need a minimum inhibitory concentration to fully protect the shrimp from IMNV and it must be 20% or more. Balasubramanian *et al* (2008) reported that aqueous extracts of *C. dactylon* showed protective effects against WSSV in *P. monodon* and the strongest antiviral activity is found at 100 mg/kg of animal body weight (Howlader *et al*., 2020). This might have been due to the method of administration employed. Since, shrimp is a slow feeding animal and has a short intestine, the absorption ability in shrimp must be lower than in human beings. Accordingly, only a small amount of extract will reach the target. The binding gel encapsulated pellet feed coated with plant extract prevents the loss of the plant extract during feeding (Balasubramanian *et al*., 2008).

## CONCLUSIONS

The results of this study show that *C. dactylon* contains strong antiviral properties with many phenolic compounds. The in vivo challenge test showed that *C. dactylon* also has anti-IMNV properties that protect shrimp from IMNV infections. Application of 20% of *C. dactylon* (Feed C) has potent activity against IMNV in *P. vannamei*. Further works on isolation, characterization, and purification of this plant’s active compounds should be carried out to apply them in the shrimp culture industry.

## REFERENCES

Balasubramanian, G, Sarathi, M., Venkatesan, C., Thomas, J., Hameed, A.S.S. 2008. Fish & Shellfish Immunology Studies on the immunomodulatory effect of extract of Cyanodon dactylon in shrimp, Penaeus monodon, and its efficacy to protect the shrimp from white spot syndrome virus (WSSV). Fish and Shellfish Immunology 25:820–828. https://doi.org/10.1016/j.fsi.2008.09.002

Balasubramanian, G., Sarathi, M., Venkatesan, C., Thomas, J., Sahul Hameed, A.S. 2008. Oral administration of antiviral plant extract of Cynodon dactylon on a large scale production against White spot syndrome virus (WSSV) in Penaeus monodon. Aquaculture 279:2–5. https://doi.org/10.1016/j.aquaculture.2008.03.052

Howlader, P., Ghosh, A.K., Islam, S.S., Bir, J., Banu, G.R. 2020. Antiviral activity of Cynodon dactylon on white spot syndrome virus (WSSV)-infected shrimp: an attempt to mitigate risk in shrimp farming. Aquaculture International. https://doi.org/10.1007/s10499-020-00553-w

Kaleeswaran, B., Ilavenil, S., Ravikumar, S. 2011. Fish & Shell fi sh Immunology Dietary supplementation with Cynodon dactylon (L.) enhances innate immunity and disease resistance of Indian major carp, Catla catla (Ham.). Fish and Shellfish Immunology 31:953–962. https://doi.org/10.1016/j.fsi.2011.08.013

Jha, R. K., Haig Babikian, Y., Yousef Babikian, H., Daniel Wisoyo, S., Asih, Y., Srisombat, S., Jiaravanon, B. 2016. Effectiveness of Natural Herbal Oil Formulation against White Spot Syndrome Virus in Penaeus vannamei. J Pharmacogn Nat Prod 2:1000123. https://doi.org/10.4172/2472-0992.1000123

Nur, Y.L. 2007. INFECTIOUS MYONECROSIS VIRUS (IMNV) IN PACIFIC WHITE SHRIMP, Litopenaeus vannamei IN INDONESIA 139–146.

Prasad, K.P., Shyam, K.U., Banu, H., Jeena, K., Krishnan, R. 2017. Infectious Myonecrosis Virus (IMNV) – An alarming viral pathogen to Penaeid shrimps. Aquaculture 477:99–105. https://doi.org/10.1016/j.aquaculture.2016.12.021

Senapin, S., Phewsaiya, K., Briggs, M., Flegel, T.W. 2007. Outbreaks of infectious myonecrosis virus (IMNV) in Indonesia confirmed by genome sequencing and use of an alternative RT-PCR detection method. Aquaculture 266:32–38. https://doi.org/10.1016/j.aquaculture.2007.02.026

